# A Computational Model of Coarctation of the Aorta in Rabbits: Ventricular and Ascending Aortic Remodeling

**DOI:** 10.1101/2023.10.30.564452

**Authors:** Ashley A. Hiebing, Matthew A. Culver, John F. LaDisa, Colleen M. Witzenburg

## Abstract

Coarctation of the aorta (CoA) is a common congenital cardiovascular lesion that typically presents as a localized narrowing of the proximal descending thoracic aorta just distal to the left subclavian artery. While improvements in surgical and catheter-based techniques have increased short-term survival, there is a high long-term risk of hypertension after CoA correction and a reduced average lifespan despite treatment. Computational models can be used to estimate ventricular and arterial remodeling, potentially serving as key tools in developing a mechanistic understanding of the interplay between pre-correction hemodynamics, post-correction recovery, and long-term hypertension risk. In this study, we developed a lumped parameter model of the heart and circulation to simulate aortic coarctation. We then used the model to estimate changes in ventricular thickness and ascending aortic compliance from imaging and catheterization data collected in rabbits with untreated and corrected CoA that used the current putative clinical treatment threshold (≥20 mmHg). Model outputs were compared to reported stroke volume, ejection fraction, systolic and diastolic ascending aortic pressures, peak ascending aortic flow, mean and peak aortic blood pressure gradients, and upper-to-lower body flow split, with all results falling within one standard deviation of the data for control, untreated CoA, and corrected CoA groups. In the untreated CoA and corrected simulations, a decrease in ascending aortic compliance was necessary to match reported hemodynamics, suggesting the rabbits exposed to CoA ≥20 mmHg underwent vascular remodeling that persisted after repair.

## 1 INTRODUCTION

Coarctation of the aorta (CoA) is a congenital cardiovascular lesion that typically presents as a localized narrowing of the proximal descending aorta just distal to the left subclavian artery. Each year, 1 in every ∼1,800 infants born in the United States is diagnosed with CoA, making it one of the most common congenital heart defects (Mai et al. 2019). While surgical or catheter-based interventions can restore flow and reduce the blood pressure gradient across the coarctation site to normal levels, successful repair of CoA does not ensure long-term cardiovascular health. Even after CoA correction, structural and functional changes in the heart and vasculature frequently occur, leading to long-term complications such as re-coarctation, aortic aneurysms, and hypertension (Cohen et al. 1989; Kenny et al. 2011; Choudhary et al. 2015; Egbe et al. 2021). In particular, despite early and effective surgical repair, the incidence of hypertension in treated CoA patients has been reported to range from 30-50% (O’Sullivan 2002; Kenny et al. 2011; Sendzikaite et al. 2020, 2022).

Maladaptive ventricular and aortic remodeling have been implicated in much of the morbidity presenting after treatment for CoA (Xu et al. 1997; Ou et al. 2008; Menon et al. 2012a). Prior to correction, patients with severe or long-standing CoA typically exhibit high systemic pressure and above-normal left ventricular (LV) mass. However, following successful CoA repair, the relationship between hypertension and LV hypertrophy is unclear. Though some studies report an association between late-onset hypertension and LV hypertrophy (Bocelli et al. 2013; Rinnström et al. 2016; Ağbaş et al. 2020), others indicate LV hypertrophy is equally likely in normotensive and hypertensive CoA patients post-correction (Krogmann et al. 1993; Eerola et al. 2007; Sendzikaite et al. 2020). After repair, antecedent LV hypertrophy begins to regress. In many patients, however, LV hypertrophy resumes following this regression. This pattern has been reported in both the presence and absence of hypertension in adult and pediatric patients (Eerola et al. 2007; Egbe et al. 2021).

In the proximal aorta, histologic analysis of tissue from untreated coarctation patients indicates increased stiffness, presenting as fibrosis, intimal thickening, and collagen degradation (Dunnill 1959; Sehested et al. 1982). This remodeling and its resultant decrease in proximal aortic compliance is strongly correlated with hypertension (Dernellis and Panaretou 2005; Mitchell et al. 2007; Kaess et al. 2012). Unfortunately, it is very challenging to assess aortic remodeling non-invasively. The two most common non-invasive metrics of aortic stiffness are pulse wave velocity and aortic distensibility. Using cardiac MRI, pulse wave velocity can be determined for a section of the aorta as the ratio of aortic length to the time delay in flow (Dijkema et al. 2018). A larger pulse wave velocity indicates a faster response and greater aortic stiffness, but results are inherently impacted by the temporal resolution of phase-contrast MRI. Aortic distensibility is the ratio of the change in aortic diameter to the change in pressure during the cardiac cycle, normalized to the minimum dimension. Therefore, aortic distensibility provides a local measure of aortic stiffness, but it requires pressure measurements. These pressures cannot be captured from the aorta noninvasively and are thus approximated from brachial cuff pressures, which can lead to an overestimation of central aortic pressure (Pauca et al. 1992). Control and normotensive corrected CoA patients have similar pulse wave velocities and distensibility in the ascending aorta, but hypertensive CoA patients exhibit significantly higher pulse wave velocity and lower distensibility (Dijkema et al. 2018).

No single clinical metric can evaluate the interplay between ventricular and aortic remodeling and cardiac and vascular function. However, computational modeling has emerged as a valuable investigative tool to study these interactions. In the context of CoA to date, modelers have used computational fluid dynamics techniques from non-invasive imaging to estimate pressure and the blood pressure gradient, wall shear stress and oscillatory shear index, and turbulent kinetic energy (LaDisa et al. 2011; Arzani et al. 2012; Wendell et al. 2013; Lee et al. 2014; Zhu et al. 2018; Ghorbannia et al. 2022). Furthermore, these approaches have been used to investigate the effects of surgical approach, stent design, and valve morphology (LaDisa, et al. 2010; Coogan et al. 2011; Jr. LaDisa et al. 2011; Wendell et al. 2013; Kwon et al. 2014). Modern fluid-structure interaction approaches can even account for aortic wall motion and elasticity, thereby permitting estimation of strain, wall tension and wall stress that have been shown to serve as stimuli for remodeling (LaDisa et al. 2011; Taelman et al. 2016; Saitta et al. 2019; Camarda et al. 2022; Azarnoosh et al. 2023). While these sophisticated finite element methods are able to capture geometric detail of the aortic arch, there has been a resurgence of interest in lumped parameter approaches (Ralovich et al. 2015; Shi et al. 2019), as they enable coupling of the aorta to the heart and circulation, provide realistic boundary conditions, and are often computationally inexpensive. Lumped parameter models have been successfully used in isolation to model aortic coarctation, demonstrating increased simulation speed, reduced parameter requirements, and the ability to inherently estimate aortic compliance (Keshavarz-Motamed et al. 2011, 2015). Computer models could be key tools to developing a mechanistic understanding of the interplay between pre-correction and post-correction hemodynamics to long-term remodeling and the potential risk for hypertension. Thus, the objective of this investigation was to 1) develop a lumped parameter model of the heart and circulation capable of simulating aortic coarctation, and 2) utilize the model to estimate changes in ventricular thickness and ascending aortic compliance from imaging and catheterization data collected using a rabbit model of CoA and correction. Analysis of the model fitting scheme and parameter uncertainty, as well as comparison to histologic observations were then conducted to evaluate model results.

## 2 METHODS

A previously published predictive computational model of cardiac hypertrophy and hemodynamics in canines (Witzenburg and Holmes 2018) was modified to replicate the reported cardiac hypertrophy and hemodynamics of three groups of rabbits (Wendell et al. 2017): a control group, an untreated coarctation group, and a corrected coarctation group.

### 2.1 Control Group

The original canine model (Witzenburg and Holmes 2018) consists of a lumped parameter circulatory model linked to a strain-based ventricular growth function. For the current study, parameters associated with LV and ascending aortic hypertrophy and remodeling were fitted using the circulatory model alone. Simulating CoA requires delineation between the ascending and descending aorta. To that end, the systemic arterial compartment was split into ascending aorta (AA), upper body (UB) arteries, and descending aorta (DA) compartments, as shown in Figure 1. Thus, the final model was composed of eight compartments: LV, RV, systemic veins (SV), pulmonary veins (PV), AA, UB, DA, and pulmonary arteries (PA). Briefly, the ventricles were simulated using time-varying elastances, pulmonary arterial behavior was simulated by a three-element Windkessel, distal vessels were represented by capacitors in parallel with resistors, and valves were represented by pressure-sensitive diodes.

**Fig. 1.**
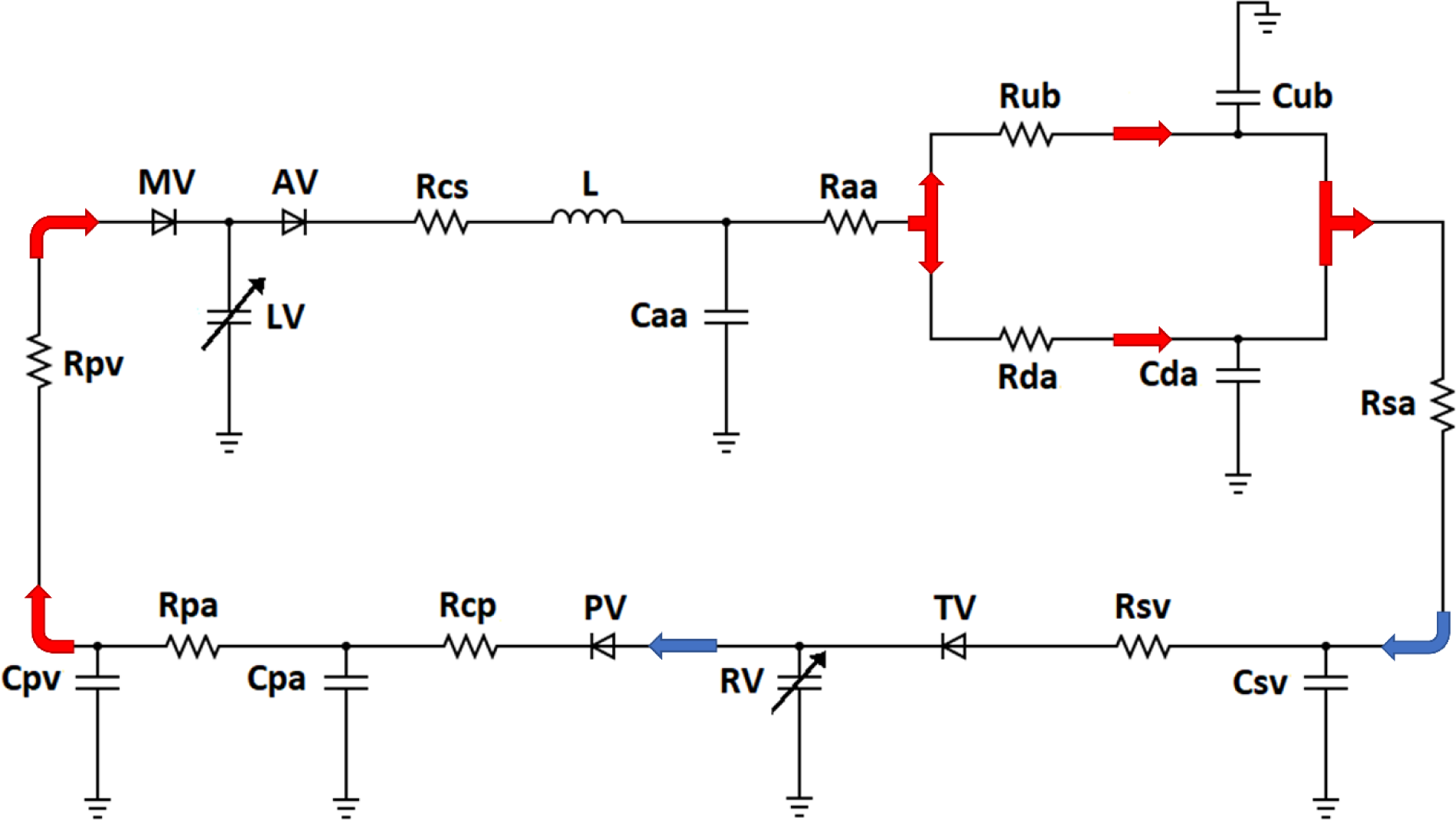
Schematic of the circuit model used to simulate the pressure-volume relationship of the cardiovascular system. The left and right ventricles (LV and RV) were simulated using time-varying elastances. Ascending aorta behavior was simulated by a four-element Windkessel consisting of a characteristic resistance (Rcs), a capacitance (Caa), an inertance (L) and an arterial resistance (Raa). The pulmonary arterial behavior was simulated by a three-element Windkessel (Rcp, Cpa, Rpa). Other vessels were represented by capacitors in parallel with resistors for the upper body arteries (Cub and Rub), descending aorta (Cda and Rda), systemic veins (Csv and Rsv), and pulmonary veins (Cpv and Rpv). Lower body systemic arteries were represented with a resistance (Rsa). Pressure sensitive diodes (TV, PV, MV, and AV) represented the tricuspid, pulmonary, mitral, and aortic valves, respectively. Arrows indicate the direction of blood flow and blood oxygenation levels

Model parameters were allometrically scaled from the original canine values assuming a rabbit weight of 3.47 kg (Wendell et al. 2017) and a canine weight of 23.4 kg (Witzenburg and Holmes 2018, 2019). Then, parameters governing the AA, UB, and DA compartments, along with stressed blood volume and ventricular parameters Ees and B (described in detail below) were tuned using the built-in MATLAB function *fminsearch* based on data from Wendell et al. (Wendell et al. 2017). Mean-squared error between reported control and model stroke volume, ejection fraction, systolic and diastolic ascending aortic pressures, peak ascending aortic flow, and mean and peak aortic blood pressure gradients (Wendell et al. 2017), as well as upper to lower body flow split (unpublished data) were minimized. Heart rate was inputted directly into the model. Input parameters are listed in Table 1.

**Table 1:** Parameter Values for Eight-Compartment Control Rabbit Model.

| **Table 1: Parameter Values for Eight-Compartment Control Rabbit Model** |  |  |
| --- | --- | --- |
| **Heart Parameters** |  |  |
| **Parameter** | **LV** | **RV** |
| End-systolic Elastance (Ees), mmHg/mL | 67.6 | 15.8 |
| Unloaded Volume (V0), mL | 0.19 | 0.19 |
| Coefficient for EDPVR (B), mmHg | 0.006 | 0.006 |
| Coefficient for EDPVR (A), mL<sup>-1</sup> | 2.44 | 2.44 |
| Ventricular wall volume (Vwall), cm<sup>3</sup> | 3.210 | - |
| **Circulation Parameters** |  |  |
| Heart Rate (HR), beats/min | 156<sup>+</sup> |  |
| Stressed Blood Volume (SBV), mL | 45.6 |  |
|  | **Pulmonic** | **Systemic** |
| Characteristic Resistance (Rcp/Rcs), mmHg·s·mL<sup>-1</sup> | 0.251* | 0.475 |
| Ascending Aortic Resistance (Raa), mmHg·s·mL<sup>-1</sup> | - | 0.696 |
| Upper Body Vascular Resistance (Rub), mmHg·s·mL<sup>-1</sup> | - | 23.1 |
| Descending Aortic Resistance (Rda), mmHg·s·mL<sup>-1</sup> | - | 0.608 |
| Vascular Resistance (Rpa/Rsa), mmHg·s·mL<sup>-1</sup> | 1.26* | 15.3 |
| Venous Resistance (Rpv/Rsv), mmHg·s·mL<sup>-1</sup> | 0.063* | 0.063* |
| Ascending Aortic Capacitance (Caa), mL/mmHg | - | 0.087 |
| Upper Body Vascular Capacitance (Cub), mL/mmHg | - | 0.002 |
| Descending Aortic Capacitance (Cda), mL/mmHg | - | 0.292 |
| Venous Capacitance (Cpv/Csv), mL/mmHg | 0.445* | 2.52* |
| Pulmonary Vascular Capacitance (Cpa), mL/mmHg | 0.297* | - |
| Characteristic Aortic Impedance (L) | - | 4.35E-05 |

<sup>+</sup>directly input from (Wendell et al. 2017)
\*allometrically scaled from canine parameters (Witzenburg and Holmes 2018), assuming a canine weight of 23.4 kg and a rabbit weight of 3.47 kg.
EDPVR, end-diastolic pressure-volume relationship

Ventricular end-systolic and end-diastolic pressure-volume relationships were defined by

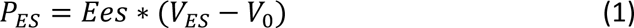

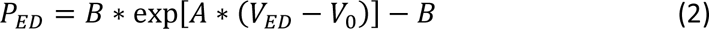

respectively, where *Ees* was end-systolic elastance, *V0* was the unloaded volume, and *A* and *B* were coefficients describing the exponential shape of the end-diastolic pressure-volume relationship. Unloaded volume *V0* was calculated as follows:

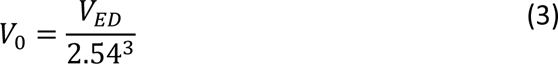

where *VED*, the end-diastolic volume, was determined by dividing reported stroke volume by reported ejection fraction (Wendell et al. 2017). From this, *A* was calculated by rearranging Equation 2:

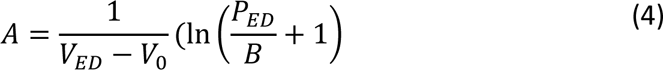

where *PED*, the end-diastolic pressure, was set to 7 mmHg (Mahaffey et al. 1995). Right ventricular parameters were kept at the same values as the left, with the exception of end-systolic elastance, which was set at 23% of the LV value (Burkhoff and Tyberg 1993).

To determine the ventricular parameters for the untreated coarctation and corrected groups, values for control LV dimensions and material properties were required. Assuming a spherical ventricle, the unloaded radius, *r0*, was

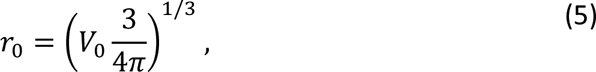

and unloaded thickness, *h0*, was

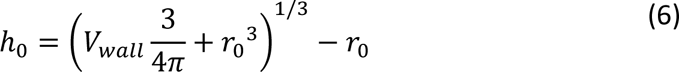

where *Vwall* was the ventricular wall volume. *Vwall* was set such that the model end-diastolic thickness matched the control group mean anteroseptal end-diastolic wall thickness reported by Wendell et al. (Wendell et al. 2017). Hoop stress relates the control unloaded ventricular dimensions (*r0* and *h0*) and pressure-volume relationship parameters (*A*, *B*, *V0*, and *Ees*) to the intrinsic ventricular material property parameters (*a*, *b*, and *e*) via the following relationships (Witzenburg and Holmes 2018):

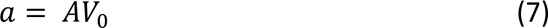

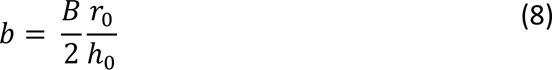

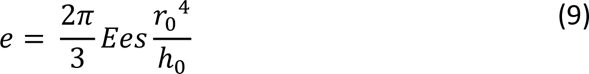

Next, the circulatory and ventricular parameters were used to calculate pressures and volumes in each compartment as well as the flows between them. The model was run until it reached steady state, defined as compartmental volumes at the beginning and end of the cardiac cycle being within 0.05 mL of each other. Total aortic flow was determined by integrating the flow from the LV to the AA over the cardiac cycle. Hemodynamic outputs (stroke volume, ejection fraction, systolic and diastolic ascending aortic pressures, peak ascending aortic flow, mean and peak aortic blood pressure gradients, and upper to lower body flow split) were saved and compared to measurements, with success defined as model outputs falling within one standard deviation of the data.

### 2.2 Coarctation Group

Aortic coarctation was simulated in three steps. First, CoA is characterized by a stenosis in the DA, therefore the DA resistance (Rda) was increased and the compliance (Cda) was decreased. Second, following prolonged CoA, ventricular hypertrophy is observed, therefore ventricular thickness was increased. Third, CoA results in remodeling of the proximal aorta involving increased thickness with reduced smooth muscle function and elastin fragmentation (Menon et al. 2012a), therefore the characteristic aortic resistance (Rcs) and AA resistance (Raa) were increased and the AA capacitance (Caa) was decreased. Parameter changes are listed in Table 2.

**Table 2:** Parameter adjustments for each group.

| Table 2: Parameter adjustments for each group |  |  |  |
| --- | --- | --- | --- |
| Parameter | Control | CoA | Corrected |
| $R_{da}$ | 0.608 | 2.536 | 0.579 |
| $C_{da}$ | 0.292 | 0.019 | 0.289 |
| $V_{wall}$ | 3.210 | 7.065 | 5.393 |
| $R_{aa}$ | 0.696 | 4.75 | 1.07 |
| $R_{cs}$ | 0.475 | 1.025 | 0.543 |
| $C_{aa}$ | 0.087 | 0.043 | 0.051 |

Hypertrophy was assumed to be concentric, therefore the coarctation unloaded radius, *c_r0*, and unloaded volume, *c_V0*, were kept at control values. Pathology was assumed to only alter ventricular thickness, therefore ventricular material properties (*a*, *b*, and *e*) were also kept at control values. Thus, of the remaining coarctation ventricular parameters,

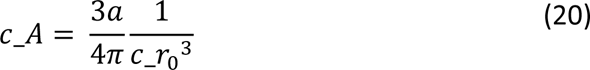

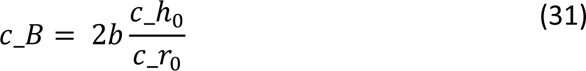

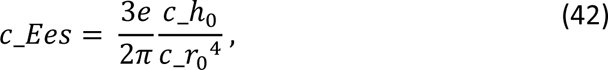

changes in unloaded thickness linearly affected only passive stiffness parameter *c_B* and elastance parameter *c_Ees*. As in the control group, *Vwall* was set to match the reported mean end-diastolic anteroseptal wall thickness. Results were compared with the reported hemodynamics of the coarctation group. Rda, Cda, Rcs, Raa, and Caa were optimized by minimizing the error in stroke volume, ejection fraction, systolic and diastolic ascending aortic pressures and flow, mean and peak aortic blood pressure gradients, and upper to lower body flow split.

### 2.3 Corrected Group

Corrected CoA was simulated much like the coarctation group. However, to model the correction of aortic coarctation, Rda and Cda were reset to control values. Despite successful correction, similar levels of proximal aortic remodeling (thickening, reduced smooth muscle function, and elastic fragmentation) have been observed long after CoA repair, suggesting some extent of proximal aortic remodeling is irreversible (Xu et al. 1997; Vogt et al. 2005; Menon et al. 2012a). Therefore, as in the coarctation simulation, Rcs and Raa were increased and Caa was decreased. *Vwall* was again set to match the reported mean end-diastolic anteroseptal wall thickness in the corrected group.

## 3 RESULTS

### 3.1 Parameter Fitting

Out of 23 total input parameters, 12 were fitted to produce hemodynamic outputs matching reported measurements in the control group. Of the remaining 11 parameters, 8 were allometrically scaled from their respective canine values (Witzenburg and Holmes 2018, 2019). Unloaded LV cavity volume, *V0*, was calculated from the reported control group stroke volume and ejection fraction (Equation 3). Heart rate was inputted directly and *Vwall* was set to 3210 mm^3^ to produce the end-diastolic anteroseptal thickness of 2.44 mm reported by Wendell et al. (Wendell et al. 2017) Model outputs for the control simulation matched reported measured stroke volume, ejection fraction, systolic and diastolic ascending aortic pressures, peak ascending aortic flow, mean and peak blood pressure gradients (Wendell et al. 2017), and upper to lower body flow split (unpublished data) within 0.1 standard deviations (Table 3).

**Table 3:** Model results compared to reported experimental measurements.

|  |  |  |  |  |  |  |  |  |  |
| --- | --- | --- | --- | --- | --- | --- | --- | --- | --- |
| Index | Control | Model | Z-score | CoA | Model | Z-score | Corrected | Model | Z-score |
| BPG, mean (mmHg) | 3.27 ± 4.32 | 3.59 | 0.08 | 20.1 ± 4.75 | 19.4 | 0.15 | 2.61 ± 3.46 | 4.48 | 0.54 |
| BPG, peak (mmHg) | 10.8 ± 7.34 | 10.3 | 0.07 | 31.0 ± 8.64 | 31.5 | 0.05 | 17.0 ± 4.32 | 14.6 | 0.55 |
| SV (mL) | 1.87 ± 0.53 | 1.84 | 0.06 | 1.99 ± 0.56 | 1.92 | 0.13 | 1.93 ± 0.26 | 1.92 | 0.04 |
| EF % | 60.6 ± 4.82 | 60.6 | 0 | 66.4 ± 6.24 | 62.5 | 0.63 | 65.0 ± 7.88 | 65.9 | 0.12 |
| Upper/lower flow split | 0.30 ± 0.04 | 0.30 | 0 | 0.37 ± 0.04 | 0.34 | 0.64 | 0.33 ± 0.04 | 0.30 | 0.54 |
| Maximum AA flow (mL/min) | 1435 ± 110 | 1435 | 0 | 1445 ± 143 | 1413 | 0.23 | 1704 ± 68.3 | 1704 | 0 |
| AA SBP (mmHg) | 67 ± 7.67 | 67 | 0 | 99 ± 17.99 | 93 | 0.35 | 69 ± 8.73 | 71 | 0.25 |
| AA DBP (mmHg) | 56 ± 9.79 | 56 | 0 | 74 ± 25.93 | 65 | 0.35 | 54 ± 10.05 | 54 | 0 |

As discussed previously in Section 2.2 and illustrated in Table 2, several parameters needed to be adjusted from their fitted control values to match measured hemodynamics in the untreated CoA rabbits. To simulate the observed increases in mean and peak blood pressure gradients (BPG) across the coarctation, Rda was increased 4.2x and Cda was decreased 15x. The mean CoA group anteroseptal wall thickness reported by Wendell et al. (Wendell et al. 2017) of 4.42 mm was matched by setting *Vwall* 7065 mm^3^. To match AA pressures and flow, Raa was increased 6.8x, Rcs was increased 2.2x, and Caa was decreased 2.0x.

In the corrected group, the diameter of the coarctation was largely restored after the aortic ligature dissolved. Therefore, Rda and Cda were set to control values. *Vwall* was set to 5393 mm^3^ to produce the reported mean end-diastolic anteroseptal thickness of 3.70 mm. Although there was a trend toward larger measurement than the reported thickness in the control group, the difference was not statistically significant. Ascending aortic parameters Raa, Caa, and Rcs required fitting to match hemodynamic outputs, though at smaller magnitudes than in the CoA simulation: Raa was increased 1.5x, Caa was decreased 1.7x, and Rcs was increased 1.1x.

### 3.2 Group Comparison

The untreated CoA simulation produced a large increase in ventricular pressure, as expected, along with a slight increase in stroke volume compared to the control group. In the corrected simulation, LV pressure was largely restored to control values, as expected, and there was a minor reduction in end-systolic and end-diastolic volumes (Figure 2).

**Fig. 2.**
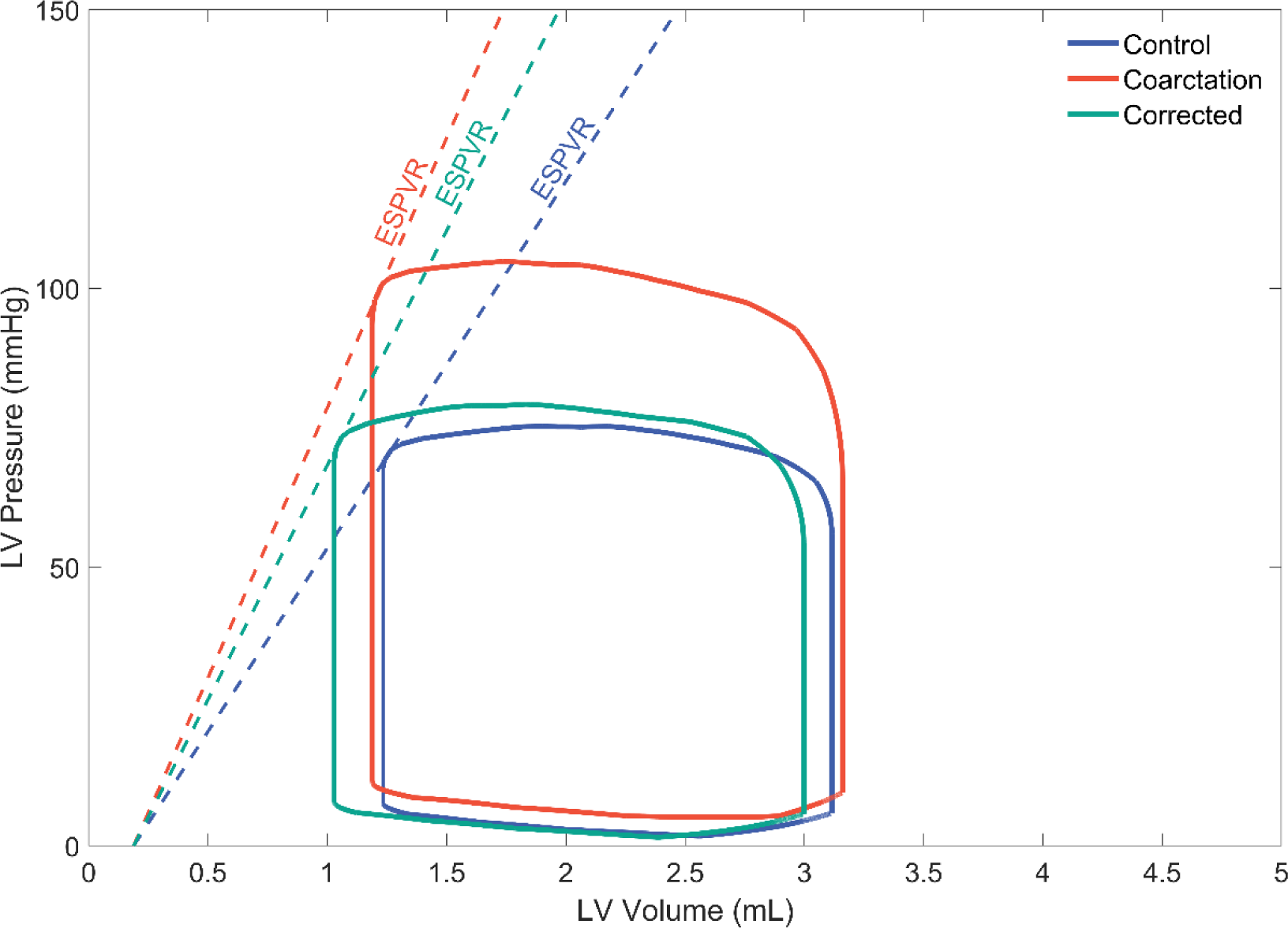
Simulated pressure-volume loops for the control (blue), coarctation (red), and corrected (green) groups. Dashed lines represent end systolic pressure volume relationships (ESPVR)

Ascending aortic systolic and diastolic blood pressures matched reported measurements well, falling within half a standard deviation for all groups (Figure 3). The blood pressure gradients across the coarctation region also agreed well with reported measurements, falling within two-thirds of a standard deviation for all groups. Stroke volume was within one-sixth of a standard deviation, and ejection fraction was within three-fourths of a standard deviation. The ratio of upper body to lower body flow was within one standard deviation for all groups. Maximum aortic flow was within a quarter of a standard deviation for all three groups. (Figure 4). Complete model results for all groups are listed in Table 3.

**Fig. 3.**
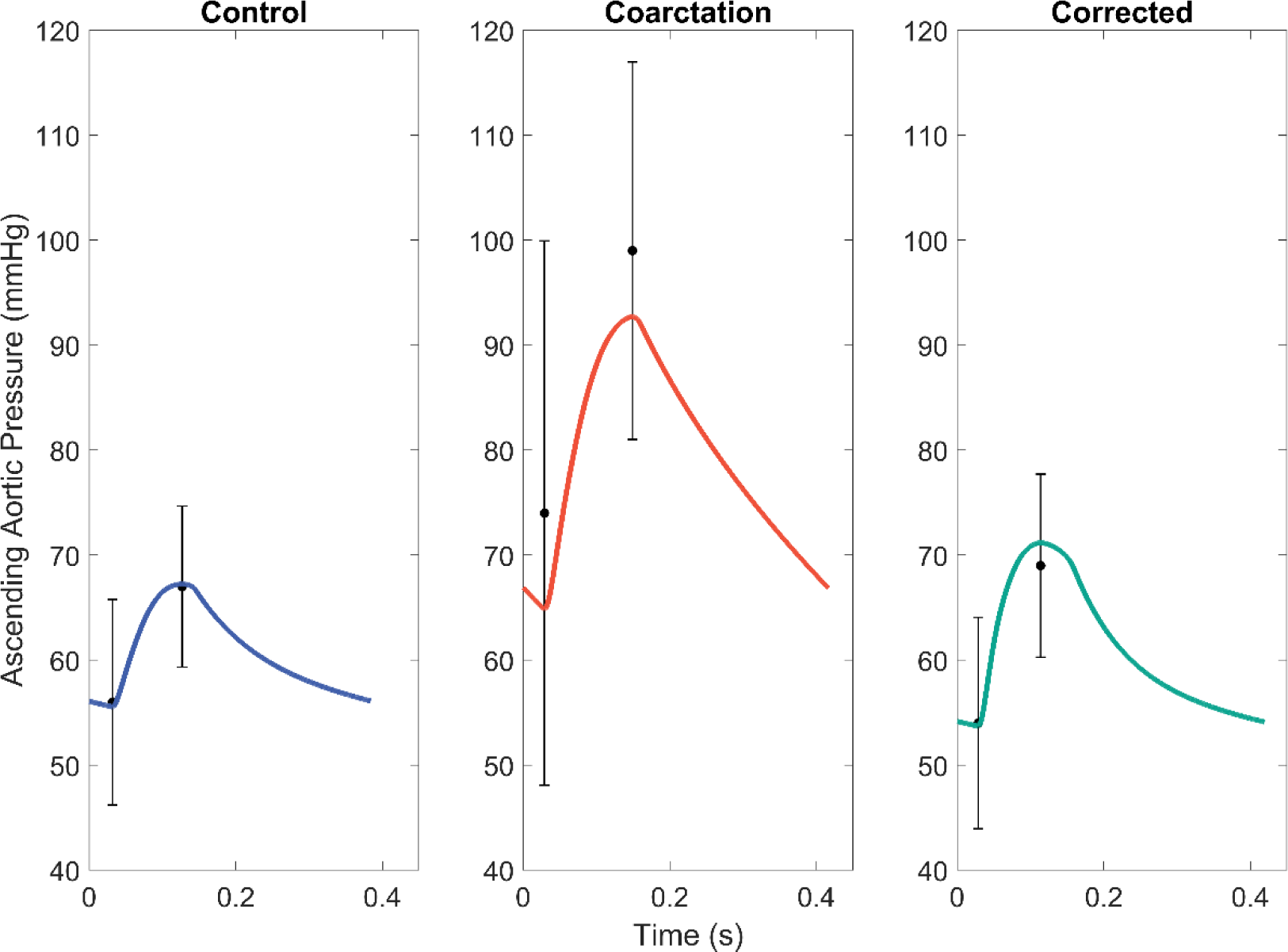
Model ascending aortic systolic and diastolic pressures over one cardiac cycle for the control (blue), coarctation (red), and corrected (green) groups, compared to measured data (black) *(Wendell et al. 2017)*. Error bars indicate one standard deviation.

**Fig. 4.**
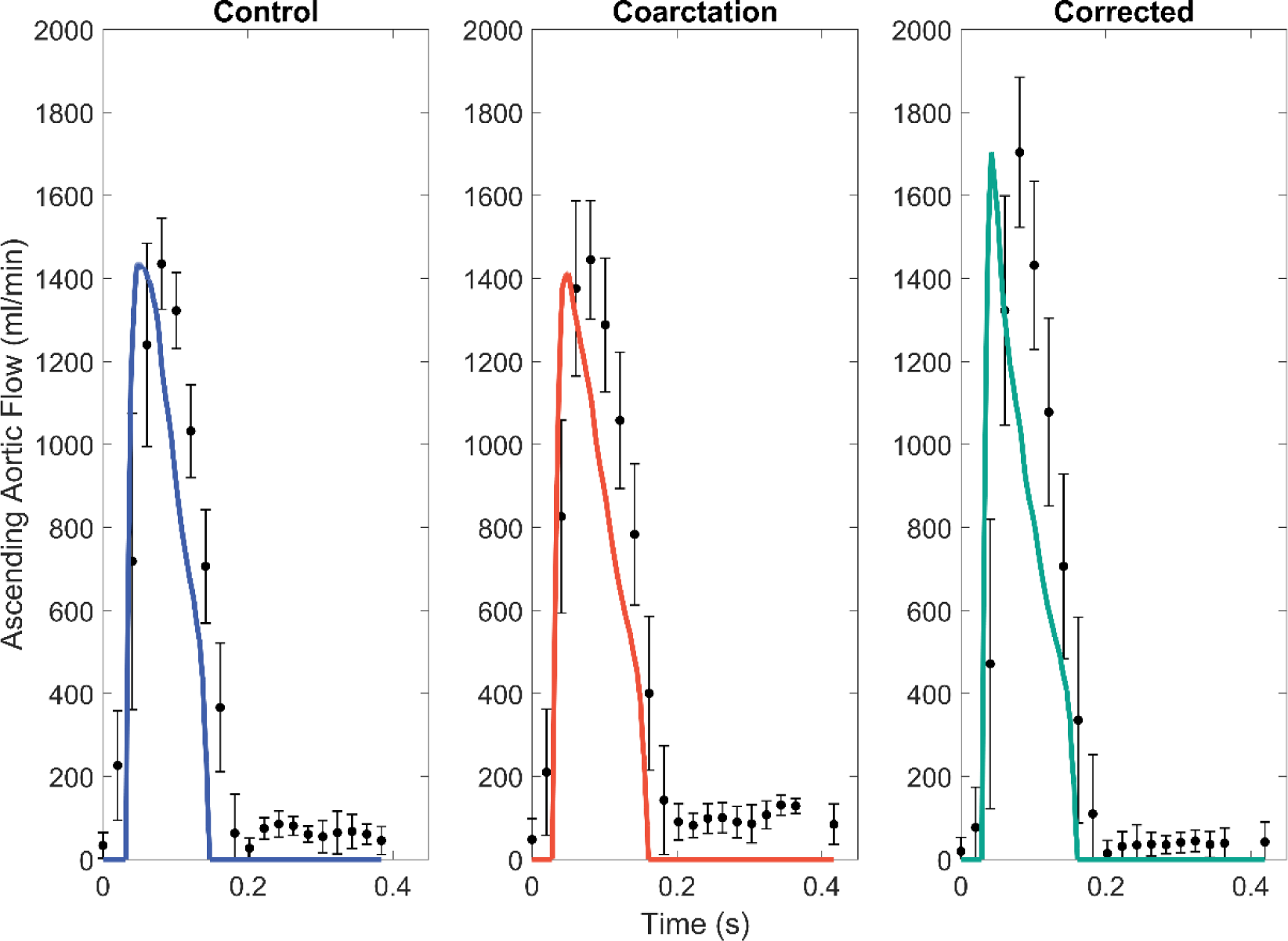
Model ascending aortic flow over one cardiac cycle for the control (blue), coarctation (red), and corrected (green) groups, compared to measured data (black) *(Wendell et al. 2017)*. Error bars indicate one standard deviation.

## 4 DISCUSSION

Previous studies of CoA have shown an increase in the stiffness of the ascending aorta before and after corrective surgery when compared to controls (Xu et al. 1997; Vogt et al. 2005; Menon et al. 2012a). In particular, histologic evidence of increases in ascending aortic thickness without increases in functional elastin or smooth muscle have been reported for both the CoA and corrected groups in the rabbit model (Menon et al. 2012b, a). These findings were replicated from the non-terminal data (Wendell et al. 2017) by our computer model: in both the untreated CoA and corrected simulations, a decrease in ascending aortic compliance was necessary to match reported hemodynamics, suggesting the rabbits with CoA underwent vascular remodeling that persisted after repair.

### 4.1 Persistent LV and Proximal Aortic Remodeling

In Wendell et al.’s study (Wendell et al. 2017), the corrected group experienced restored diameter of the coarctation region after dissolving of the Vicryl suture used to create the coarctation, which resulted in a reduced blood pressure gradient when compared to the untreated CoA group. Despite this reduction, the corrected group experienced a larger peak aortic flow than the control or CoA groups. However, total flow and ascending aortic pressures did not differ from control, indicating flow acceleration in the corrected group and suggesting irreversible aortic stiffening proximal to the coarctation. We explored this by modeling three parameter cases for the CoA and corrected simulations (Figures 5 and 6, respectively).

**Fig. 5.**
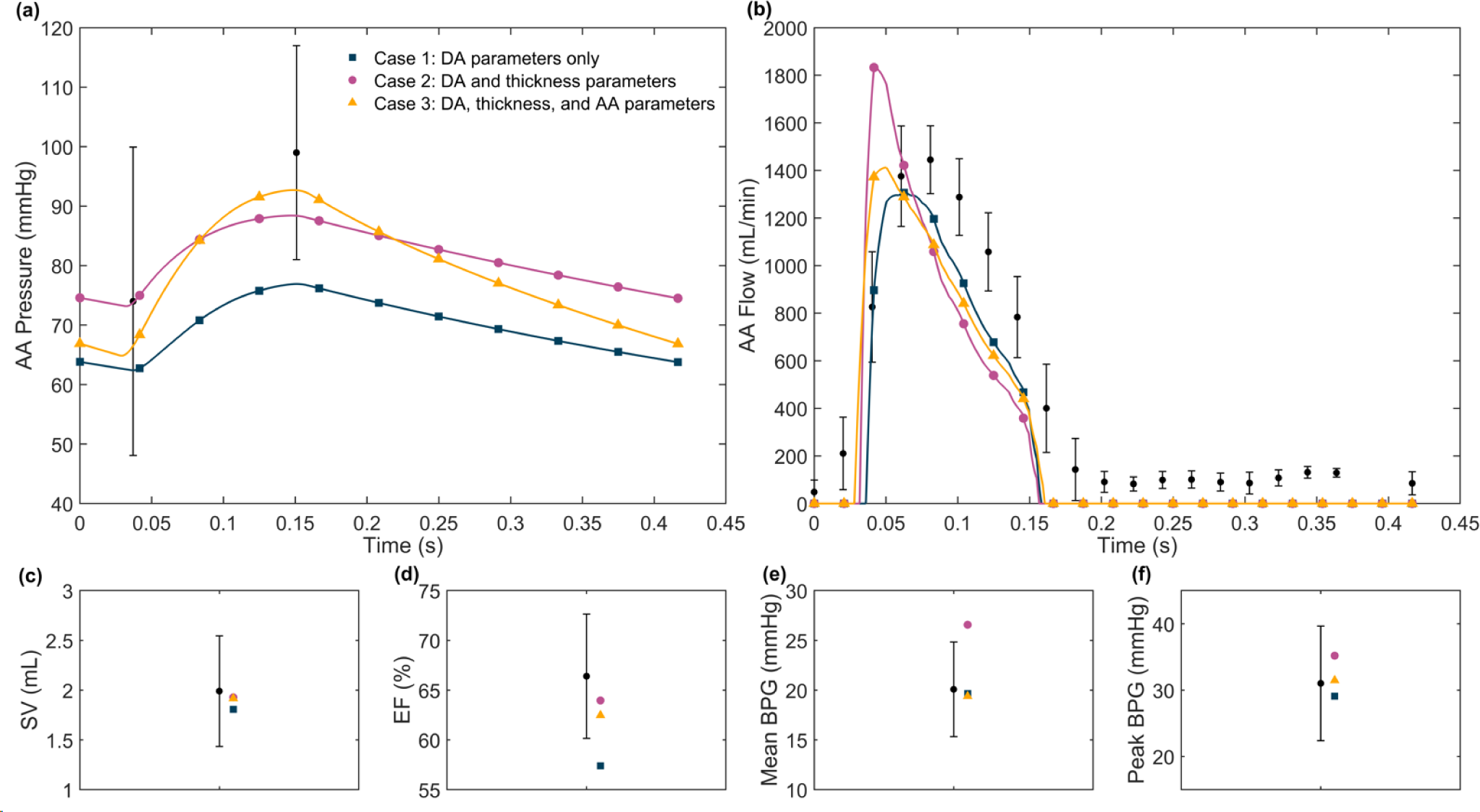
Parameter cases for the untreated coarctation simulation, with model outputs compared to reported measurements for **(A)** ascending aortic pressure, **(B)** ascending aortic flow, **(C)** stroke volume, **(D)** ejection fraction, **(E)** mean blood pressure gradient, and **(F)** peak blood pressure gradient *(Wendell et al. 2017)*. In **Case 1** (blue squares), only the descending aorta parameters (Rda and Cda) were allowed to change. For **Case 2** (purple dots), Rda and Cda were allowed to change, and Vwall was set to produce a loaded end-diastolic thickness equivalent to measured untreated CoA group mean anteroseptal wall thickness *(Wendell et al. 2017)*. In **Case 3** (yellow triangles), thickness was kept at the same value as Case 2, and Rda, Cda and ascending aorta parameters (Raa, Caa, and Rcs) were allowed to change. All other parameters were kept at control values. Error bars indicate one standard deviation

**Fig. 6.**
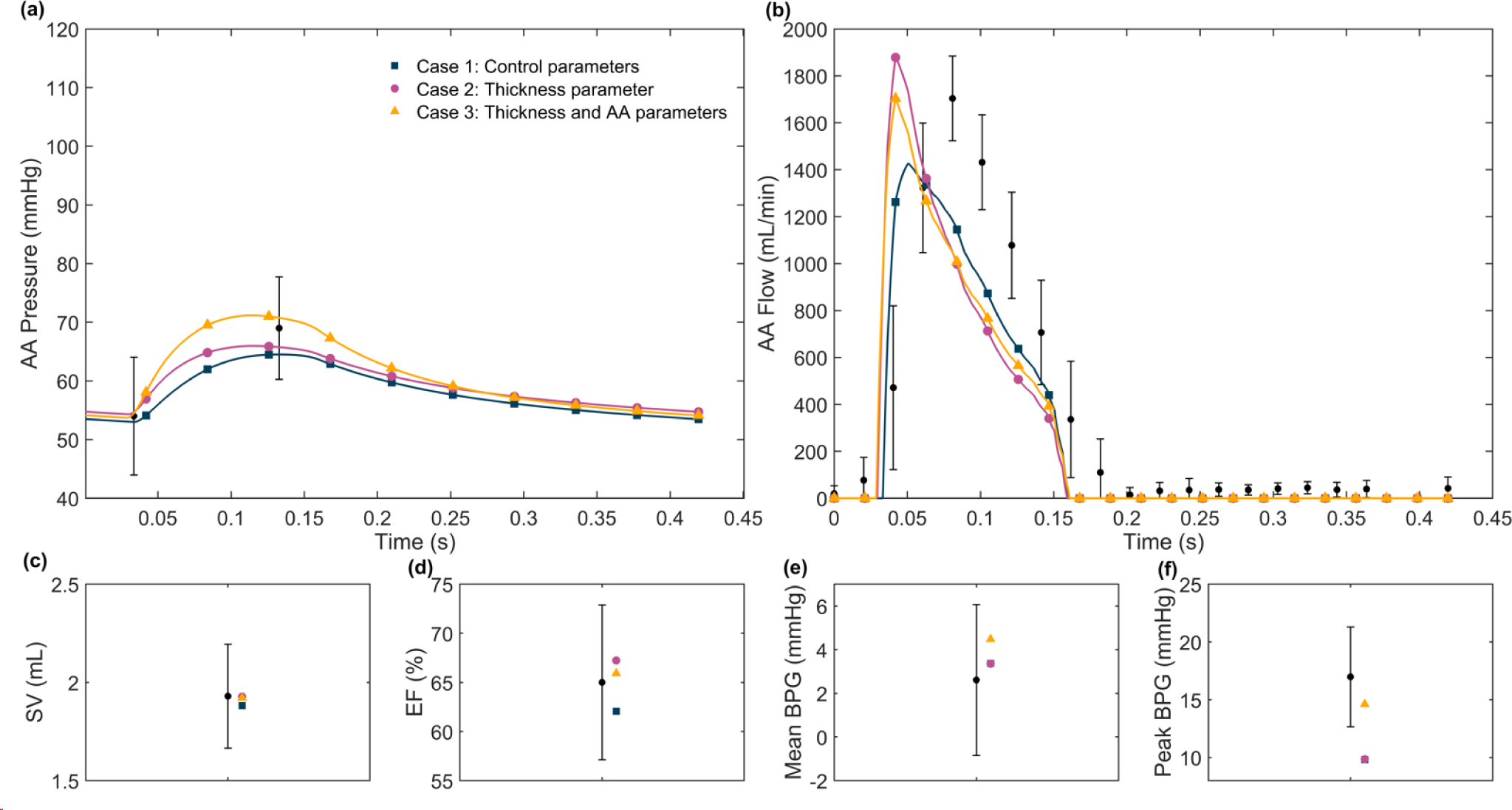
Parameter cases for the corrected simulation, with model outputs compared to reported measurements for **(A)** ascending aortic pressure, **(B)** ascending aortic flow, **(C)** stroke volume, **(D)** ejection fraction, **(E)** mean blood pressure gradient, and **(F)** peak blood pressure gradient *(Wendell et al. 2017)*. In **Case 1** (blue squares), all parameters were kept at control values. For **Case 2** (purple dots), Vwall was set to produce an end-diastolic thickness equivalent to measured corrected group mean anteroseptal wall thickness *(Wendell et al. 2017)*. In **Case 3** (yellow triangles), thickness was kept at the same value as Case 2, and ascending aorta parameters (Raa, Caa, and Rcs) were allowed to change. Outputs for Cases 1 and 2 overlap in (E) and (F). All other parameters were kept at control values. Error bars indicate one standard deviation

### CoA group

- **Case 1:** For the first case, only descending aortic parameters (Rda and Cda) were optimized, leaving thickness and ascending aortic parameters equivalent to control values. Though these parameters were sufficient to match stroke volume, diastolic ascending aortic pressure, and mean and peak blood pressure gradient measurements, they were insufficient to match ejection fraction, peak ascending aortic flow, and systolic ascending aortic blood pressure.
- **Case 2:** In the second case, Vwall was set to produce an end-diastolic thickness equivalent to the mean anteroseptal wall thickness measured in the untreated CoA group. Rda and Cda were allowed to change, as in Case 1. The increase in thickness produced an ejection fraction and ascending aortic diastolic blood pressure matching measured data. However, peak aortic flow and the mean blood pressure gradient were too high.
- **Case 3:** For the third case, Vwall, Rda, and Cda were kept at their Case 2 values. In addition, ascending aortic parameters Raa, Rcs, and Caa were allowed to change. With the inclusion of these parameters, peak aortic flow and the mean blood pressure gradient were reduced to the appropriate values.

### Corrected group

- **Case 1:** To represent a complete reversal of the coarctation, the first case for the corrected group used the same parameters as the control group. Though these parameters produced a stroke volume, ejection fraction, mean blood pressure gradient, and ascending aortic systolic and diastolic pressure within one standard deviation of measured data, peak aortic flow and peak blood pressure gradient were too low.
- **Case 2:** In the second case, Vwall was set to produce an end-diastolic thickness equivalent to the mean anteroseptal wall thickness measured in the corrected group. All other parameters were kept at control values. With the increase in thickness, peak aortic flow output was within a standard deviation of measured data. However, the peak blood pressure gradient remained below the measured value.
- **Case 3:** For the third case, Vwall was set to the same value in Case 2, and ascending aortic parameters Raa, Rcs, and Caa were allowed to change. Only by altering these parameters could the peak blood pressure gradient be matched.

To further evaluate our modeling results, we used a frequentist analysis (Smith 2014; Colebank et al. 2021) to estimate the uncertainty in the parameters associated with ventricular and ascending aortic remodeling: Vwall, Raa, Rcs, and Caa. This approach considers the local sensitivity of each best-fit parameter to experimental data and the sum-squared error produced by best-fit model predictions, enabling computation of parameter confidence intervals. Figure 7 shows the 95% confidence internals for the control model as well as the best-fit untreated CoA and corrected models. Caa confidence intervals for both the untreated CoA and corrected models were well below the control model, further confirming that a reduction in ascending aortic compliance was necessary to capture the changes in the experimental data between both the control and untreated CoA rabbits and the control and corrected rabbits. In addition, the Vwall confidence intervals did not overlap, indicating a larger increase in ventricular muscle mass from control was necessary to match the untreated CoA data as opposed to the corrected data. The control and corrected Raa confidence intervals almost overlapped, potentially implicating ascending aortic stiffening or thickening as the mechanism of reduction in Caa, rather than a reduction in lumen radius.

**Fig. 7.**
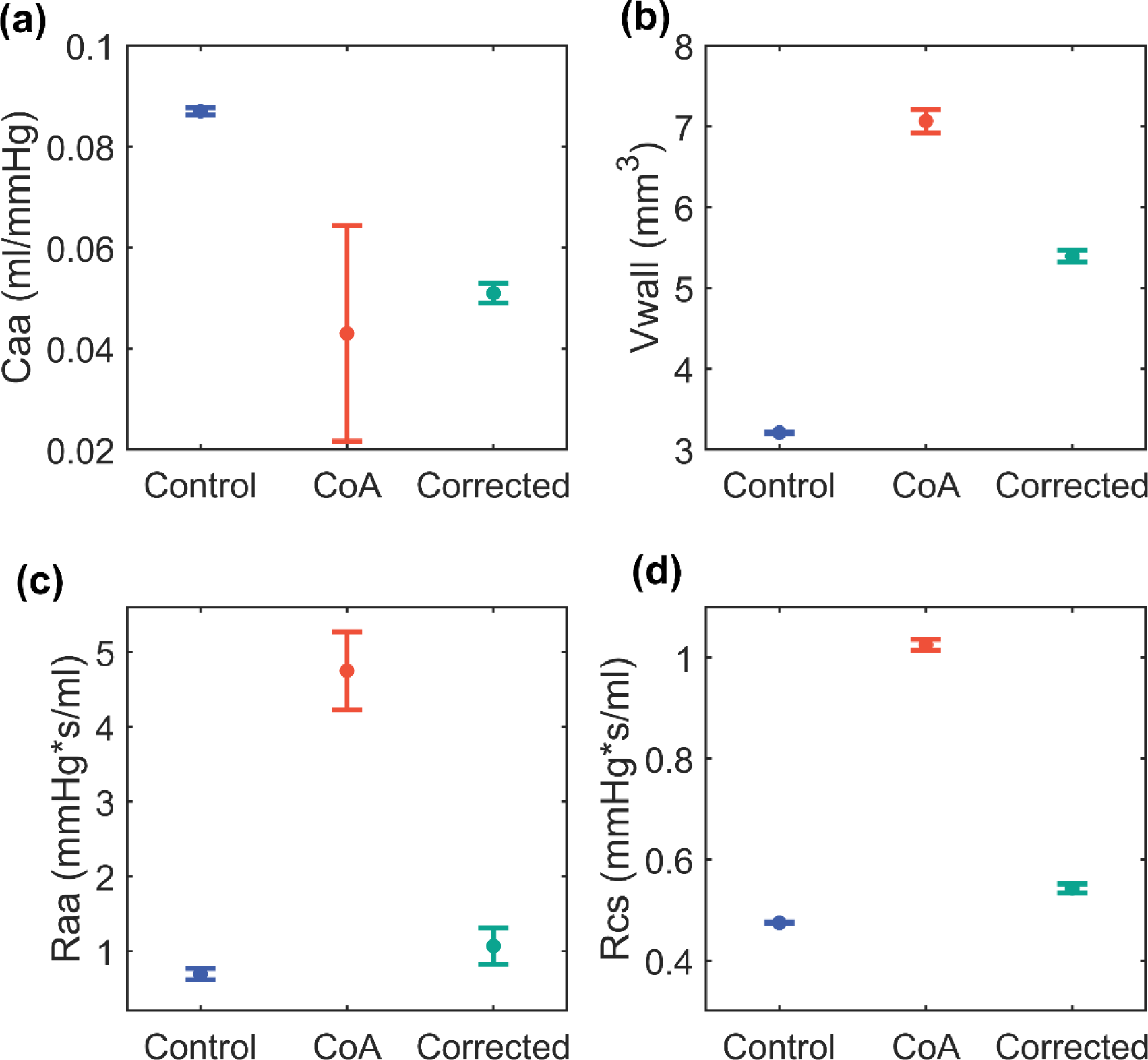
Best-fit model parameters for the control, CoA, and corrected simulations associated with ventricular and ascending aortic remodeling, including the model parameter indicating **(A)** the ascending aortic capacitance (Caa), **(B)** the volume of myocardium (Vwall), **(C)** the ascending aortic resistance (Raa), and **(D)** the systemic characteristic resistance (Rcs). Error bars indicate 95% confidence intervals determined for each parameter from a frequentist uncertainty analysis

We also performed additional simulations to determine if the hemodynamic changes in the corrected group could be explained by incomplete repair of the coarctation. Allowing descending aortic parameters to change while keeping other parameters equivalent to control values failed to replicate the increase in peak aortic flow measured in the corrected group. As in the untreated coarctation group, simulating persistent remodeling in both the left ventricle and the ascending aorta was necessary to match reported hemodynamics.

### 4.2 Limitations

We assumed a simplified spherical geometry when representing the ventricles in our model. Though this assumption did not impact our ability to match reported hemodynamics or ventricular dimensions within a standard deviation, a more realistic geometry would produce a physically meaningful ventricular wall volume and may facilitate a closer model fit to experimental data. The simple nature of this model does make it extremely fast, however, requiring 1.4 seconds to run on a desktop computer with 16 GB RAM, a 64-bit operating system, and a 3.0 GHz Intel Core i7-9700 CPU, expediting both automated parameter fitting and uncertainty analysis.

Collateral vessels can form to attenuate the hemodynamic consequences of the coarctation (Steffens et al. 1994; Fox et al. 2019). While collateral vessels have not been directly observed in this animal model (Menon et al. 2012b, a; Ghorbannia et al. 2022), the caliber of intercostal arteries at harvest indicate it is likely they were present but not possible to resolve by MRI. Experimentally, Azarnoosh et al. (Azarnoosh et al. 2023) measured the total upper body flow in this rabbit model by adding flows for the arteries proximal to the coarctation (the right and left coronary and subclavian arteries) and the total systemic flow by integrating the area under the ascending aorta PC-MRI waveform. Thus, while collateralization was not explicitly considered in the computer model, the increase in upper body flow and reduction in blood pressure gradient that can result from collateralization were considered during model fitting.

Changes in arterial compliance are represented in our computer model by changing the value of aortic capacitors Caa and Cda. These capacitance changes cannot differentiate between compliance changes due to geometry (changes in wall thickness or radius) and stiffness (due to microstructural changes in elastin, collagen, and smooth muscle). Terminal histologic analysis of ascending aortic tissue indicates these effects are likely occurring simultaneously. Both untreated CoA and corrected rabbits exhibited increases in ascending aortic thickness and reductions in functional elastin and smooth muscle (Menon et al. 2012b, a). Multiscale microstructural (Witzenburg et al. 2017; Rolf-Pissarczyk et al. 2021) and continuum constitutive (Holzapfel et al. 2004) computer models can capture the effects of tissue microstructure on overall mechanical behavior. While they would clearly be capable of delineating the effects of microstructural remodeling and vessel thickness, fitting model parameters would necessitate terminal structural and geometric data. Predictive computational models (Alford et al. 2008; Tsamis et al. 2009; Cyron and Humphrey 2014; Latorre et al. 2019; Mahutga and Barocas 2020), which simulate alterations in tissue constituent mass due to pathologic loading, could potentially address this concern but have not yet been applied to aortic coarctation.

Lastly, we assumed constant myocardial material properties in all three groups. Animal studies of LV pressure overload caused by aortic banding have shown ventricular remodeling includes collagenous fibrosis (Perlini et al. 2005). Additionally, histologic analysis indicates patients with heart failure and a preserved ejection fraction alongside hypertension have higher myocardial collagen content than healthy controls or those without hypertension (Zile et al. 2015). However, biopsy is not feasible in most patients. Late gadolinium enhanced cardiovascular MRI is a common non-invasive approach to quantify focal or replacement myocardial fibrosis, such as that seen in ischemic heart disease, and T1 mapping is emerging as a newer technique capable of detecting diffuse fibrosis, which is more common in hypertensive heart disease. While both techniques have been applied to CoA patients post-correction, fibrosis was not detected (Luijendijk et al. 2013; Pieper et al. 2019). We were able to conduct preliminary histologic analyses on left ventricular samples from one control and one untreated CoA rabbit from the replicated experimental study. Short axis slices were stained with picrosirius red and imaged under polarized light to visualize collagen. We did not observe any obvious difference in collagen content, and automated quantification indicated that collagen volume fraction was the same in the control and untreated CoA ventricular samples (2.6% vs. 2.5%, respectively), although this finding was not statistically confirmed. In the future, if we find evidence that myocardial material properties change with CoA or its correction, accounting for this in the computational model would be relatively straightforward.

### 4.3 Conclusion

There is increasing evidence that the degree of vascular and ventricular remodeling due to CoA is likely linked to the duration and severity of the coarctation (Menon et al. 2012a; Azarnoosh et al. 2023). Though post-correction reversal of this remodeling may be incomplete or impossible, comprehensive non-terminal approaches are necessary to estimate its course and degree. Computational models of CoA could explore the relationship between duration and/or severity of CoA with remodeling, potentially becoming a valuable clinical tool in patient assessment and surgical planning. Future simulations will model rabbits with CoA of varying intensity and corrections at different timepoints. From this, we can investigate the trade-off in severity and duration of CoA to evaluate their individual roles in the timecourse and reversibility of ventricular and vascular remodeling.

